# A simple high efficiency and low cost *in vitro* growth system for phenotypic characterization and seed propagation of *Arabidopsis thaliana*

**DOI:** 10.1101/2020.08.23.263491

**Authors:** Taras Pasternak, Benedetto Ruperti, Klaus Palme

## Abstract

**Background:** Arabidopsis research relies extensively on the use of *in vitro* growth for phenotypic analysis of the seedlings and characterization of plant responses to intrinsic and extrinsic cues. For this purpose, stress-free optimal growth conditions should be set up and used as a reference especially in studies aimed at characterizing the plant responses to abiotic and biotic stresses. Currently used standard *in vitro* protocols for growth and characterization of *Arabidopsis thaliana* plants often suffer from sub-optimal composition due to an excessively high nutritional content which represents a stress per se and an experimental bias.

**Results:** We describe a simple protocol for *in vitro* growth of Arabidopsis plants in which the phenotypic analysis is based on an optimized and nutritionally balanced culture medium. We show that the protocol is robustly applicable for growth of several Arabidopsis mutants, including mutants lacking the root system. This protocol enables rapid high scale seed production *in vitro* avoiding soil usage while saving space and time. The optimized *in vitro* protocol aims at: 1) making *in vitro* growth as close as possible to natural soil conditions by optimizing nutrient balance in the medium; 2) simplifying phenotypic and molecular investigation of individual plants by standardizing all steps of plant growth; 3) enabling seeds formation also in genotypes with severe defect in the root system; 4) minimizing the amount of waste and space for plant growth by avoiding soil usage.

**Conclusions:** Here we report an optimized protocol for optimal growth of *Arabidopsis thaliana* plants to avoid biases in phenotypic observation of abiotic/biotic stress experiments. The protocol also enables the completion of the whole life cycle *in vitro* within 40-45 days and a satisfactory seed set for further propagation with no need for facilities for plant growth in soil and seed sterilisation.

## Introduction

Over the last decade, Arabidopsis has become a widely adopted model system in plant biology for a variety of reasons. These include a short life cycle, its fully annotated genome, the simplicity of the root system, the availability of natural diversity sets, large numbers of mutants and easy procedure of genetic transformation [1].

Current research largely relies on the extensive use of *in vitro* growth of seedlings for phenotypic analysis and characterization of plants’ responses to intrinsic and extrinsic cues. Physiological and molecular biology experiments, aimed at unravelling plants’ responses to abiotic and biotic stresses, in principle require starting plant material that has been grown in a nutritionally stress-free environment before and during analysis. Hence, growth conditions should as be as close as possible to the optimal. In the majority of cases researchers use culture media based on Murashige and Skoog composition [2], which has been originally designed for callus growth and is characterized by a specific nutrient balance with high nitrogen contents, a high N:P ratio (60:1.25) and with chloride as a major anion. These ratios and absolute contents do not fit with nutrient content in plant tissues and nutrient’s ratios which are widely known to be optimal for plant nutrition [3]. In many cases such a medium itself may induce stress-associated phenotypes [4]. A hallmark of this phenotype is the time-dependent change in the Arabidopsis meristem size that reaches its maximum at 5 days after germination while root growth almost stops at day 8 [5]. This impact of medium composition on plant growth and phenotypic behavior is also exampled by Arabidopsis growth on the more balanced modified Hoagland medium where the root meristem size increases up to day 12, and the root’s growth continues till 3 weeks [6, 7]. These effects are particularly relevant considering that one of the central aims of adopting *in vitro* culture of seedlings is the investigation of the root system architecture (RAS). Root architecture is notoriously dependent on nutrient availability [8, 9]. For these reasons, it is essential that for adequate interpretation of physiological and molecular results, especially when the adaptive strategies to changing environments are the focus of research, a nutrient balance as close as possible to the one existing in soil should be adopted for *in vitro* culture media.

Hydroponic culture, as an alternative for soil culture, represents a reliable procedure in terms of space and reproducibility. Hydroponics itself also requires a completely different optimal medium composition in order to avoid representing a stress per se [4].

In addition, in the commonly used protocols after initial *in vitro* growth and characterization, mutant or wild type plants are typically transferred to soil pots for further phenotypic evaluation and for propagation (seed production).

This procedure requires soil and is costly in terms of time and greenhouse space, while it generates a large amount of waste. Although the Arabidopsis time span per generation is relatively short, the completion of its life cycle from-seed-to-seed in soil requires around 6 to 8 weeks under optimal conditions in greenhouse [10]. Greenhouse conditions require significant energy inputs for plant care including watering, nutritional supplementation and protection from pests and fungi with the use of chemicals or through biocontrol. Seeds arising from greenhouse grown plants require further sterilization, a process that is not always successful and may give rise to contaminations and loss of (sometimes precious) material. Using average laboratory conditions, the full growth cycle of Arabidopsis plants usually takes up to 10 weeks and requires labor and space for successful growth and propagation. Considering these limitations, we have sought for a suitable, simple, rapid procedure to be carried out entirely *in vitro*, based on a culture system with nutrient composition optimized for plant growth and characterization. This system minimizes nutritional stress, by being more closely similar to natural conditions than the commonly used MS medium, and resembles growth of plants in greenhouse in terms of seed set and seed maturation. A procedure aimed at obtaining seed-set entirely *in vitro* and at shortening the seed-to-seed generation time in Arabidopsis, has been previously described [11]. This procedure was based on the germination of mature seeds in picloram and benzyladenine in the first generation and of immature seeds in the following generations, thus allowing the authors to obtain up to 12 generations annually [11]. However, the amount of seeds generated by using this procedure appeared limited and the method was shown to be successfully applicable only to some Arabidopsis ecotypes. In addition, immature seeds, while accelerating the propagation speed, do not allow making precise quantitative characterization of each generation and introduce a significant bias in experiments aimed at evaluating phenotypes related to all aspects of plant biology.

The method we present here allows completing the whole *Arabidopsis* life cycle within 40-45 days, starting from mature seeds while ensuring phenotypic observation of plants in the absence of external or additional perturbations (e.g. hormone treatments or use of immature seeds) and a satisfactory seed set for further investigations.

## Methods

### Plant material

*Arabidopsis thaliana* (L.) Heyhn lines Col-0, auxin response mutants *tir1* [12], *tir1xafb1,2,3* [13], *plt1,2,3* [14], *miao* [15]).

### Reagents

1. TK1 medium, for *in vitro* growth of seedlings (Table 1). Stock solutions were made by the following way: x2000 Micro salts stock. Dissolve in the following order: 25 mg Na_2_MoO_4_·2H_2_O, 800 mg H_3_BO_3_, 1800 mg MnCl_2_·4H_2_O, 20 mg CuSO_4_·5H_2_O, 20 mg CoCl_2_.6H_2_O, 180 mg ZnSO_4_·7H_2_O, 80 mg NaJ in a final volume of 100ml ddH_2_O. 400x Fe-chelate stock: Dissolve 372.2 mg Na_2_EDTA·2H_2_O and 278.5 mg FeSO_4_·7H_2_O in separate glass beakers under continuous stirring at 70°C in 20 ml ddH_2_O each. Mix the solutions and prolong heating till the solution gets a gold-yellow color. Adjust volume to 50 ml. Crucial: AM (Arabidopsis medium corresponding to ½ MS, Murashige and Skoog medium) medium has an N:P:K ratio of 60:1.25:20 which is sub-optimal for plant growth ([2]). A more balanced medium in terms of nutrient ratio (TK1 medium) (see text for explanations) is suggested for *in vitro* growth of plants, especially in studies focusing on plant nutrition and stress response. Hoagland’s (Hg) medium [16] (Table 1) supplemented with 2% sucrose for *in vitro* seed propagation. Crucial: 2% sucrose is essential for high recovery of seeds.
2. Seed sterilisation solution (20% sodium hypochlorite Roth, cat. no. 9062.3).
3. Murashige and Skoog (MS) medium (Duchefa, cat. No. M0245.0050).
4. Sucrose (Roth, cat. no. 9097.1).
5. Peptone ex casein (Roth cat. No 8986.1)
6. Sterile de-ionised water.

### Materials

1. Square 12 cm Petri plates.
2. Glass tubes (Scott Duran, 18 cm) with cotton tops. Crucial: cotton top is essential for proper aeration and seed drying. When a plastic top is used, humidity remains high and seeds do not dry.
3. Forceps, Scalpels (sterile).
4. Miracloth (Calbiochem Catalog No. 475855).
5. Tops: (Rotilabor, Kultur stopfen CTE0.1).

### Procedure

Overview of procedure steps and time required for each step.

**Table 1.** Composition of different nutrients media (in mg/l) compared in this work for *in vitro* growth of Arabidopsis seedlings (TK1) and seed propagation (Hg). The composition in terms of macro- and micro-nutrients is reported for Hoagland medium (Hg), Arabidopsis medium (AM; half-strength Murashige and Skoog Medium) and for the optimized medium described in this work (TK1).

| Compound | AM (½ MS) | TK1 | Hg |
| --- | --- | --- | --- |
| **Macronutrients** |  |  |  |
| NH<sub>4</sub>NO<sub>3</sub> | 825 | - | - |
| KNO<sub>3</sub> | 950 | 900 | 600 |
| NH<sub>4</sub>H<sub>2</sub>PO<sub>4</sub> | - | 200 | 125 |
| MgSO<sub>4</sub>·7H<sub>2</sub>O | 180 | 200 | 300 |
| KH<sub>2</sub>PO<sub>4</sub> | 68 | - | - |
| Ca(NO<sub>3</sub>)<sub>2</sub>·4H<sub>2</sub>O | - | 250 | 980 |
| CaCl<sub>2</sub>·2H<sub>2</sub>O | 220 | - | - |
| **Fe-Chelate** (2M stock) | 2.5 ml | 2 ml | 2 ml |
| **Micronutrients** |  |  |  |
| H<sub>3</sub>BO<sub>3</sub> | 3.1 | 4 | 2.86 |
| MnCl<sub>2</sub>·4H<sub>2</sub>O | 8.4 | 9 | 1.81 |
| ZnSO<sub>4</sub>·7H<sub>2</sub>O | 15 | 0.9 | 0.22 |
| Na<sub>2</sub>MoO<sub>4</sub>·2H<sub>2</sub>O | 0.125 | 0.125 | 0.12 |
| NaI | 0.38 | 0.4 | 0.75 |
| CuSO<sub>4</sub>·5H<sub>2</sub>O | 0.012 | 0.1 | 0.05 |
| CoCl<sub>2</sub>·6H<sub>2</sub>O | 0.012 | 0.1 |  |
| **Organic compounds** |  |  |  |
| Bacto-tryptone | - | 150 | 150 |
| Vitamins | ½ B5 vitamins | ½ B5 vitamins | ½ B5 vitamins |
| Sucrose | 10000 | 10000 | 20000 |
| MES | 200 | 200 | 200 |
| pH | 5.6 | 5.6 | 5.6 |

1. Seeds sterilization (can be omitted if *in vitro* formed seeds were used); approximately 50 min −1 h.
2. Plant growth on TK1 medium, for phenotypical and genetic characterization; approximately 17-22 days.
3. Transfer of plants to Hoagland medium in glass tubes; approximately 30 min.
4. Seed formation in Hoagland medium in glass tubes; approximately 7-10 days.
5. Seed drying and collection; approximately 7-10 days.

## Procedure details

### Plant growth and characterization

#### Step 1. Seed sterilization and cold treatment

Place seeds in a Miracloth (Millipore, catalog number 475855) bag and transfer to 20% NaOCl solution, containing 0,05% Tween 20. Apply short-time vacuum for 1 minute and incubate seeds for additional 7 minutes. Wash seeds in sterile distilled water 3 times, 5 minutes each, and transfer to 40 mm diameter Petri plates.

##### Comments

This step can be omitted if seeds were generated in sterile *in vitro* conditions. After sterilization, seeds are transferred to 12 cm square Petri plates containing TK1 nutrient medium in 1% agar (Carl Roth, Art. Nr.5210.2). Dishes are kept (with covers) at room temperature for 6-8 hours to allow imbibition, then they are moved to 4°C in the dark for 14-16 hours for stratification, following standard procedures.

##### Comments

The 6-8 hours imbibition at room temperature is a crucial step for much more effective synchronization of seeds germination.

#### Step 2. Seedling growth

Transfer plates to a growth room with 16/8 h day/night cycle, 24°C, with a light intensity of 80-100 µmol/m^2^/sec (or continuous light, depending on experimental strategy) for further growth. Seedlings are grown on the agar surface, as in standard AM based protocols. Roots can be collected for immuno-localization analyses at this stage, from 5-10 days old plants as described in [17]. Seedlings are kept in plates until plants transit to bolting/flowering. In wild type plants this usually takes 15-18 days starting from seed soaking.

#### Step 3. Transfer of plants to tubes with Hoagland medium for in vitro seed propagation and maturation

As soon as plants start bolting, the shoots can be transferred to tubes with Hoagland medium. The root system can be cut with a scalpel and can be collected for molecular analyses. The rosette with the flowering stem is transferred to tubes containing Hoagland medium supplemented with 0,8% agar in a sterile bench. We recommend to add in each tube not more than 2-2.5 ml of medium. The tubes are then moved back to the growth room with a 16/8 h day/night photoperiod. Seed formation and maturation occur within the next 2-3 weeks. During this time the medium in the tubes undergoes slight drying.

##### Comments

Successful seed formation can occur only if tubes are covered with cotton top and accompanied by plants drying. In the case of plastic top, water evaporation is prevented thus causing high humidity inside tubes, that prevents efficient seed formation and drying. The tops are prepared from cheesecloth (or paper towel) and cotton and can be re-used several times. Tops can be prepared from double layers of towel or cheesecloth filled with cotton and finally closed with a rope (Figure 1 and 3 A and B). Alternatively, tops can be ordered (Rotilabo®-Kulturstopfen CTE 0.1).

**Figure 1.**
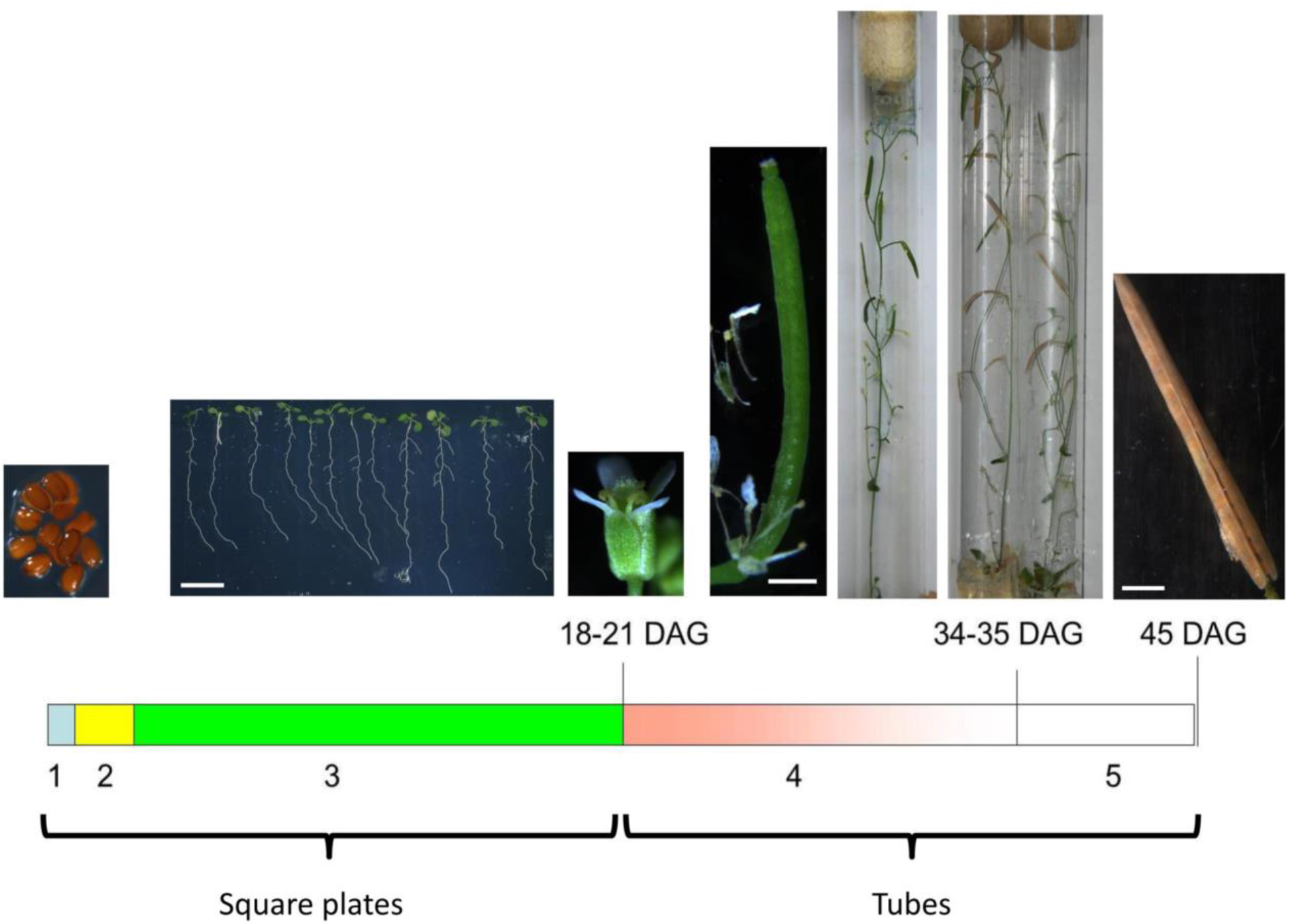
Workflow of the protocol for *in vitro* seeds formation of Arabidopsis plants.

We recommend to autoclave tubes and medium separately and add 2-2.5 ml of the medium to the tubes just before usage since medium dries after prolonged storage even at 4°C.

If a low amount of the medium remains in the tubes while the siliques and seeds are still green, we recommend to add 1 ml of fresh liquid Hoagland’s medium to the tube and extend cultivation for 1-2 more weeks.

#### Step 4. Seed collection

After seeds become completely dry, they can be collected in sterile Petri plates (40 mm diameter). Collect seeds in a sterile bench with sterile forceps directly into sterile plates or tubes. In this case the seed sterilization step can be omitted.

A workflow for the whole procedure of *in vitro* growth of Arabidopsis plants for seeds formation is shown in Figure 1.

## Results and discussion

### Effects of media composition on plant growth

Plant phenotyping and molecular analysis is an important task in plant biology. Such analysis should be performed in conditions which are as close as possible to the natural soil conditions because plant phenotypic responses generate through a complex interaction of genotype with environmental conditions [7, 8, 9]. Nutrient solutions which reproduce as closely as possible the natural conditions in soils have been described previously and have been intensively used to study plant phenotypic responses to nutrient availability [6, 9]. A sub-optimal composition of the plant culture medium, potentially causing a nutritional stress, may significantly alter plant development and phenotypic characterization, since plant nutrition is a key factor in determining plant development through a complex interaction with genetic cues. Being the organs devoted to cope with nutrient availability, roots are especially sensitive to such conditions so that an alteration in root system architecture (RSA) can be easily caused by improper nutrient availabilities [9]. As an example, it was recently shown [18] that nutrient balance is a key regulator of CLE peptide signaling and, through this, of the whole root architecture. The increasing interest in characterizing the plants’ responses to abiotic stresses further poses the need for media allowing reliable analysis in automated phenotypic screening systems specifically aimed at reproducing in the lab nutritional conditions which may be closer to the plant physiological requirements in the field and do not represent a nutritional stress *per se*. This is of particular interest when the genetic factors for response to nutrient availability and homeostasis are sought for. So far the most used nutrient medium for *in vitro* growth of Arabidopsis seedlings is half strength Murashige and Skoog (MS) medium (1962) [2], generally termed AM (Arabidopsis Medium). MS medium was originally designed starting from White’s media [19] and optimized for rapid growth of tobacco callus cells in the presence of external supply of hormones [2]. This medium is characterized by a very high nitrogen content, high N:P:K ratio of 60:1.25:20 and contains chloride as a major anion (Table 1). However, the optimal N:P:K ratio for plant growth in soil and in hydroponic culture (5:1:3) is far from that found in the AM medium [3]. Moreover, chloride content should fall within micronutrient levels [20]. In addition, the concentrations of essential micro-nutrients such as Cu and Co are rather low in AM medium. Co, for example, which is an important inhibitor of ethylene production [21], is frequently overlooked as a medium component, particularly in the case of limited aeration *in vitro.* Nutritional stress may also cause epigenetic changes [22] or alterations in root morphology [23]. Thus a culture medium which strictly follows an optimal nutrient ratio and optimal concentrations of macronutrients and micronutrients would be highly desirable in order to avoid nutritional stress and allow phenotypic evaluations in conditions which are closer to real nutrient availability in the field. To this end, we suggest the adoption of a balanced culture medium (TK1) with an optimal ratio between nutrients (Table 1) for growth, phenotypic analysis and *in vitro* propagation of Arabidopsis, which can be successfully used also for mutants lacking the root system. This medium has been previously optimized for *in vitro* growth of barley [24] and alfalfa [25]. One of the main problems of *in vitro* tissue culture is the formation of precipitates in media after mixing all ions and sucrose [26, 27]; this frequently happens even before autoclaving. This problem can be circumvented by adding a low amount of enzymatically digested casein hydrolysate (Tryptone) (150 mg/l) to the medium. This addition completely prevents precipitations in the medium, while it may serve as a source of amino acids. The total nitrogen contents (12%) and amino acid content (3%) in the Tryptone may have limited impact on the processes studied. In this circumstance we have to mention that an average soil (natural conditions) contains a quite high amount of organic nitrogen/amino acids [28], that are, however, difficult to be standardized.

For the evaluation of the effects of composition of media on the root morphology we have compared root growth on AM (½ strength MS) medium and TK1 medium. Root growth proceeded significantly faster on TK1 medium in comparison with AM medium in wt plants (Figure 2). Even more pronounced effects were observed for the auxin response mutants *tir1* [12] and *tir1xafb1,2,3* [13], caused by excess nutrient stress by AM medium (Figure 2A and B). Interestingly, on TK1 medium the root meristem size is increased in both mutants and wt (Figure 2D), as well as post-mitotic cell elongation is enhanced (Figure 2 C,D). Similar results were observed for a number of other mutants. We investigated the root growth behavior of well-known mutants defective in cytokinin production (*ipt3ipt5ipt7* [29]) and *miao* ([15]) mutants on TK1 medium. It was shown that on balanced medium the effects of these mutations on root development were much milder than those observed on 1/2 MS medium [15], emphasizing the importance of using a nutritionally balanced medium for Arabidopsis growth (Table 2).

**Table 2.** Comparison of the *in vitro* (this article) and in soil seeds generation approaches in Arabidopsis. Asterisk (*) indicates averaged data.

|  | Standard growth in soil | *In vitro* growth |
| --- | --- | --- |
| **Parameters** |  |  |
| **Space per plant** | 20 sq.cm | 2 sq.cm |
| **Duration from seeds to seeds** | 60 -70 days | 40-45 days |
| **Amount of waste** | 300 ml (soil) | 3 ml (medium) |
| **Seeds per plant\*** | 10000 | 350 |
| **Standardization of growth** | Low | High |
| **Genetic seeds “purity”** | Moderate, isolation of plants by covering is required | 100 % pure from fully isolated plants |
| **Watering, greenhouse care, use of chemicals/pesticides** | Yes | No |
| **Mutants with defects in root development** | Very low yield, not applicable for mutants with severely impaired roots | Applicable to rootless mutants (e.g. *rml1*, *plt1,2,3*). |
| **Generation of plants/seeds containing N<sup>15</sup> labeling** | Not applicable | Applicable |
| **Seeds sterilization** | Required | Not required. Seeds recovered from *in vitro* grown plants are already sterile |

**Figure 2.**
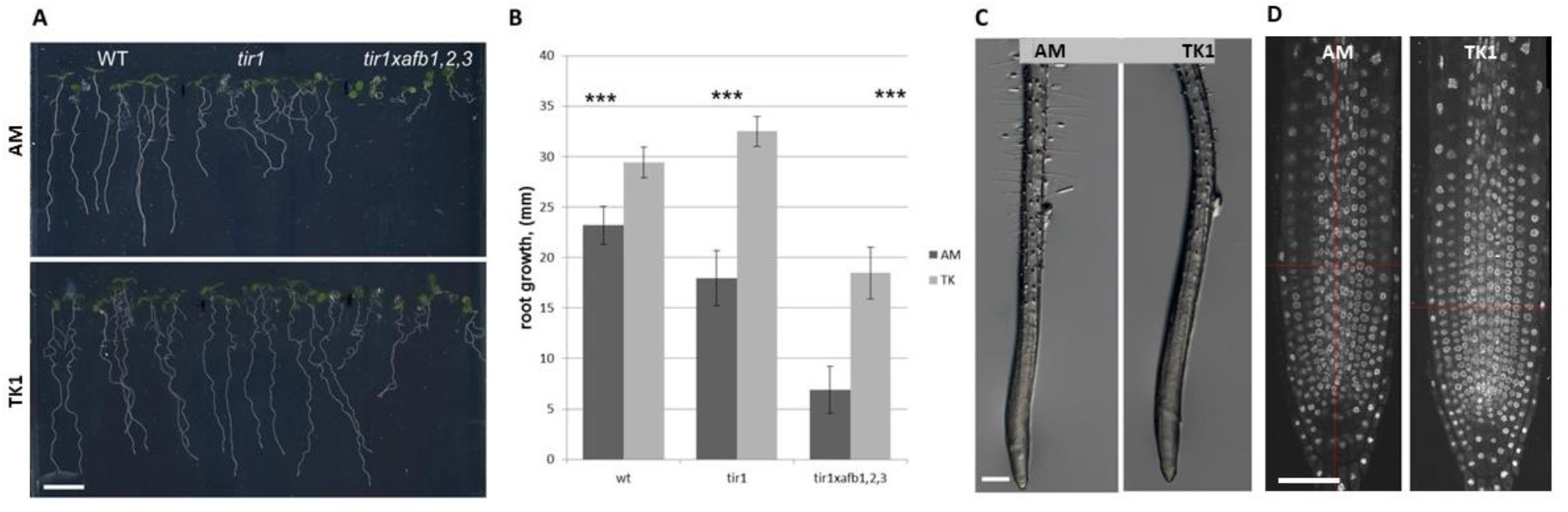
Effect of AM and TK1 medium on root development. (A) Plants of wt (Columbia), *tir1* and *tir1xafb1,2,3* mutants were cultured on AM medium for 4 days and were transferred on AM or TK1 medium thereafter. Pictures were taken after 3 additional days of growth. Scale bar - 1 cm; (B) - Quantification of root length (mm) on AM and TK1 medium. Error bar-SD; *** mean significant differences for P ≤0,001. (C) – details of root images. Scale bar - 100 µm. (D) – DAPI stained RAM. Scale bar-50 µm.

An additional important application of the protocol is that it enables the growth of plants and subsequent generation of seeds from wild type and also from mutants with severe root defects. For optimal seed formation we successfully used Hoagland medium that ensured optimal plant growth and silique formation in comparison to AM (Figure 3). As expected, seed formation requires a relatively high amount of phosphorus and lower nitrogen contents than those present in AM medium, which may prevent normal bolting and growth. Hoagland medium was developed by Hoagland and Arnon, further revised by Arnon, and has been originally designed for plant growth in hydroponic culture [21]. It provides every nutrient in optimal ratios necessary for seed formation. Presence of carbon and energy sources is crucial for seed formation; therefore addition of 2% sucrose is recommended. Both TK1 and Hoagland medium have much higher biological buffering capacity because they have a lower NH_4_/NO_3_ ratio. This ratio is important because NH_4_ uptake is much faster compared with NO_3_ uptake. This, in turn, leads to significant acidification of the medium. To avoid such acidification we suggest adding MES.

**Figure 3.**
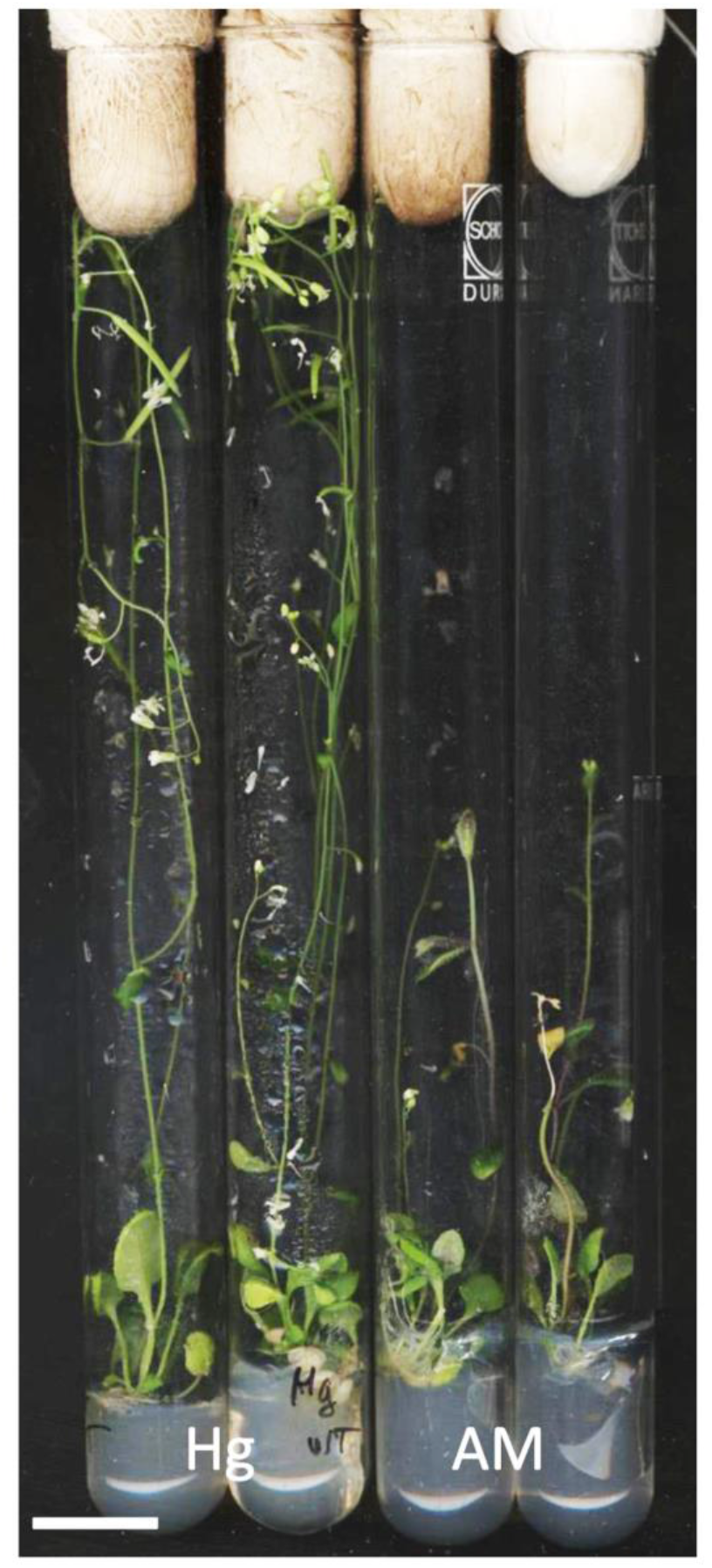
Comparison of Hg and AM medium for *in vitro* seeds reproduction. Seedlings were grown for 18 days on TK1 medium, were transferred to tubes and cultured for next 10 days.

As far as mutants with impaired root development are concerned, the *plt1,2,3* triple mutant [14], with severe defects in PIN polarity in the roots and in the development of the root system, has been used for this purpose to test the efficacy of the *in vitro* seed formation protocol. *plt1,2,3* (+/-) seeds were germinated and plants with severe defects in root formation were selected. Selected plants (Figure 4 A) were analyzed for defects in establishing PIN1 polarization in the root (Figure 4 B, C), and thereby the formation of the auxin gradient, and for a complete lack of the root system and inability to form adventitious roots, to ensure selection of homozygous seedlings. AM and Hoagland medium were compared for their performance in the process of seed formation. Plants containing small excised stems with flowers have been transferred to tubes containing either ½ MS or Hoagland medium (supplemented with 2% sucrose). After 2 weeks in the tubes strong differences were detected in the number and quality of seeds (Figure 4 D, E and F). This procedure allowed recovery of seeds from rootless mutant plants such as the *plt1,2,3*.

**Figure 4.**
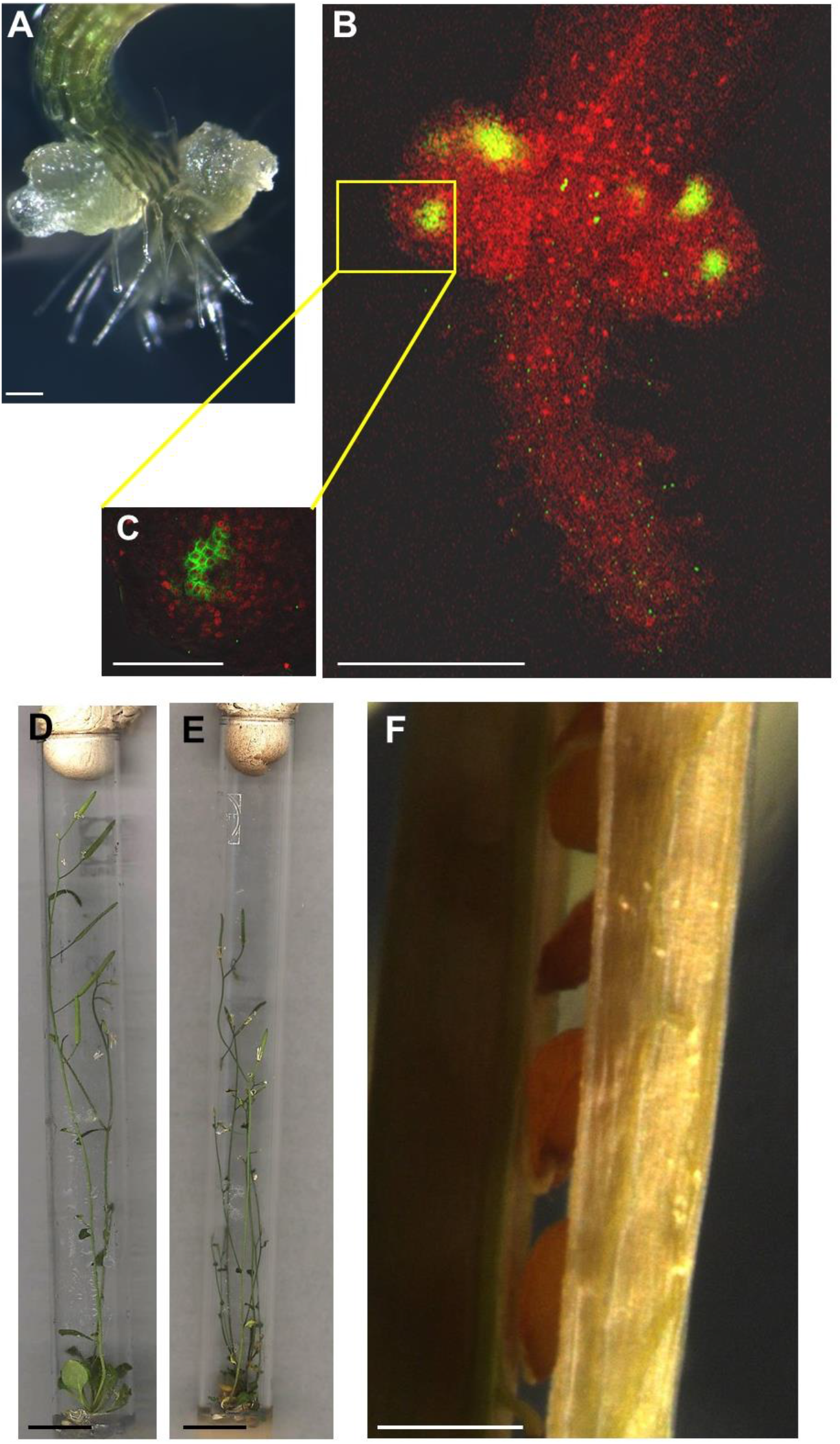
Seeds formation from homozygotic *plt1,2,3* mutant. (A) – 4 days old plt1,2,3 homozygotic seedlings. (B, C) – PIN1 immunolocalization (according to the protocol described in [2]) in the selected *plt1,2,3* mutants roots (see text for explanations). (D) and (E) - wt and *plt1,2,3* mutant plants in tubes after 35 days from seeds soaking. (F) details of a *plt1,2,3* silique at seed ripening stage. Scale bar: A, B- 100 µm; C- 50 µm; D, e – 1cm, F – 500 µm.

Overall, the application of the *in vitro* growth procedure for seed reproduction described here allows a significant reduction of time and space. The system also has been applied for generation of N15 seeds, which have been used for stable isotope labeling with amino acids in cell culture (SILAC) [30]. In our hands we were able to generate seeds which have more than 98% of N15 contents. The advantages of the procedure described here in comparison with previously published procedures are resumed in table 3.

**Table 3.**
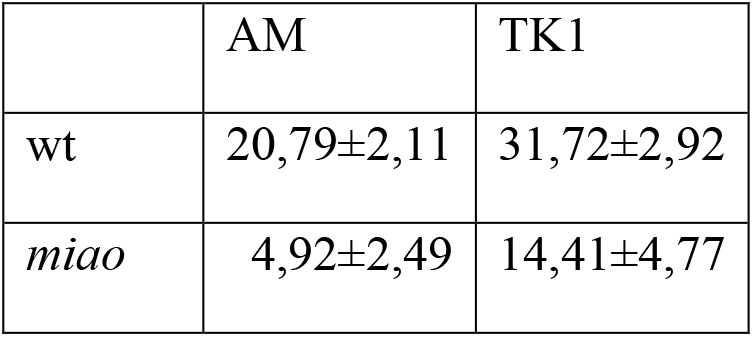
Root length of wt and *miao* mutant seedlings on AM and TK1 medium

## Conclusions

We report a simple *in vitro* system for the growth, phenotyping and seed reproduction of *Arabidopsis thaliana* plants. The system is based on two steps: 1) seedling cultivation on vertical plates on optimal medium that allows selection and characterization of the plants during the first 3 weeks of growth; 2) Shoot cultivation on modified Hoagland medium during the next 2-3 weeks which allows to generate high amounts of genetically pure and sterile seeds.

The system enables to shorten the Arabidopsis cycle from seed to seed to about 45 days, while significantly reducing the amount of waste generated from growth in soil pots and obtaining genetically pure lines by completely avoiding cross-pollination. The sterility of all steps of plant growth until seeds drying removes the need for further sterilization steps for the growth of the next generations. In wild type Arabidopsis plants we were able to generate more than 300 seeds per plant on average, while in the case of mutants with severe root impairment the amount of seeds was restricted to 100 per tube but still allowed efficient reproduction.

## Declarations

### Ethics approval and consent to participate

Not applicable.

### Consent for publication

Not applicable.

### Availability of data and materials

All data generated or analyzed during this study are included in this published article.

### Authors contributions

TP performed the experiments; TP, BR and KP interpreted and discussed the results and wrote the manuscript.

## Acknowlegements

The authors thank Katja Rapp for many stimulating ideas and excellent technical assistance.

